# The perceptual categorization of multidimensional stimuli is hierarchically organized

**DOI:** 10.1101/2021.09.29.462467

**Authors:** Chi Chen, Livia de Hoz

## Abstract

As we interact with our surroundings, we encounter the same or similar objects from different perspectives and are compelled to generalize. For example, we recognize dog barks as a distinct class of sound, despite the variety of individual barks. While we have some understanding of how generalization is done along a single stimulus dimension, such as frequency or color, natural stimuli are identifiable by a combination of dimensions. To understand perception, measuring the interaction across stimulus dimensions is essential. For example, when identifying a sound, does our brain focus on a specific dimension or a combination, such as its frequency and duration? Furthermore, does the relative relevance of each dimension reflect its contribution to the natural sensory environment? Using a 2- dimension discrimination task for mice we tested untrained generalization across several pairs of auditory dimensions in a naturalistic and automatized behavioral paradigm. We uncovered a perceptual hierarchy over the tested dimensions that was dominated by the sound’s spectral composition. This hierarchy could reflect the relevance of the different dimensions in natural stimuli and their potentially associated differential shaping of neuronal tuning. Mice could learn to pay more attention to dimensions low in the hierarchy, but this learning was more rigid and did not generalize as flexibly. Stimuli are thus not perceived as a whole but as a combination of their features, each of which weights differently on the dentification of the stimulus according to an established hierarchy.

## Introduction

Animals, including humans, learn to discriminate between the diverse sensory stimuli in their environments and to generalize their behaviors to new instances of those stimulus types. For example, rain can sound dramatically different as it falls on different surfaces and yet we have no problem classifying it as rain and discriminating it from speech. What is the basis of this ‘carving nature at its joints’ (Plato, 370BC)^1^, by which our brains separate the world into objects which fall into different classes e.g., footsteps, motorbikes, human voices, the smell of rain? Do we discriminate two stimuli to be of different classes, or generalize them into the same class, because of isolated physical properties such as frequency or color, or because of their characteristics in some more abstract multi-dimensional space? Is this classification determined by the relative sensitivity of sensory neurons to different stimulus dimensions? And is this a reflection of the contribution of these dimensions to the information in natural stimuli?

Most of our knowledge of stimulus perception is derived from tests along a single stimulus dimension. For example, psychophysics and electrophysiology studies have shown that the visual system can discriminate or categorize along color, contrast, or orientation. Or that auditory system is able to discriminate or categorize along sound frequency^2–6^, intensity^5–8^, frequency modulation direction^9–12^ or amplitude modulation frequency^4,13^. Yet stimulus discrimination along a single dimension can be influenced by stimulus features outside that dimension, as has been demonstrated in auditory processing^14–18^. While in the visual modality, dimension integration has been studied by training animals such as pigeons to use dimension combinations^19^, or by asking human subjects to identify pop-up stimuli of different level of complexity^20^, little is known about how multiple dimensions within any given modality are naturally integrated when classifying environmental stimuli. To understand the interactions between stimulus dimensions, as well as the hierarchy of importance of the dimensions, we focused on the auditory domain. We trained mice to perform 2-dimensional sound discriminations before testing untrained generalization. For example, we trained them to classify a high-frequency upward-sweep as safe and a low-frequency downward-sweep as unsafe. These two sounds differed, therefore, in both frequency and the direction of the modulation. We then tested generalization by examining how the mice classified novel stimuli that varied independently in both dimensions. For example, we determined if they judged a low-frequency upward-sweep to be safe, as the direction would suggest, or unsafe, as the frequency would suggest. By measuring how mice perceived and acted upon these changes we aimed to understand which dimensions tend to dominate in the discrimination of sounds (e.g. frequency or sweep direction).

## RESULTS

We trained mice to discriminate between two frequency-modulated sounds, with one sound being ‘safe’, indicating water would be delivered at a spout upon a nose-poke, and the other being ‘unsafe, indicating an aversive air puff would be delivered instead of water. The two sounds differed in two out of four dimensions; frequency range, sweep direction, modulation rate, and duration (Fig 1a). For example, for sounds differing in frequency range and sweep direction, the safe sound could be a 9-18 kHz upward-sweep (Fig. 1b, blue frame), and the unsafe sound, a 6-3 kHz downward-sweep (Fig. 1b, red frame). Once the animals had learned to discriminate between the safe and the unsafe sounds, we sparsely introduced novel sounds (always safe) that varied in one or both of the two dimensions. Examining whether the mice responded to these sounds as ‘safe’ or ‘unsafe’ allowed us to explore how they generalize what they had learned. For example, a novel sound could have the frequency range of the safe sound but the direction of the unsafe sound, forcing the animal to make a choice whether to treat the new sound as similar to the safe or to the unsafe (Fig. 1b, other sounds in matrix). Measuring generalization gave us access to the mouse’s perception of the two sound dimensions at play and to establish whether decisions were based on just one of the dimensions or on both dimensions (Fig. 1c).

**Fig. 1.**
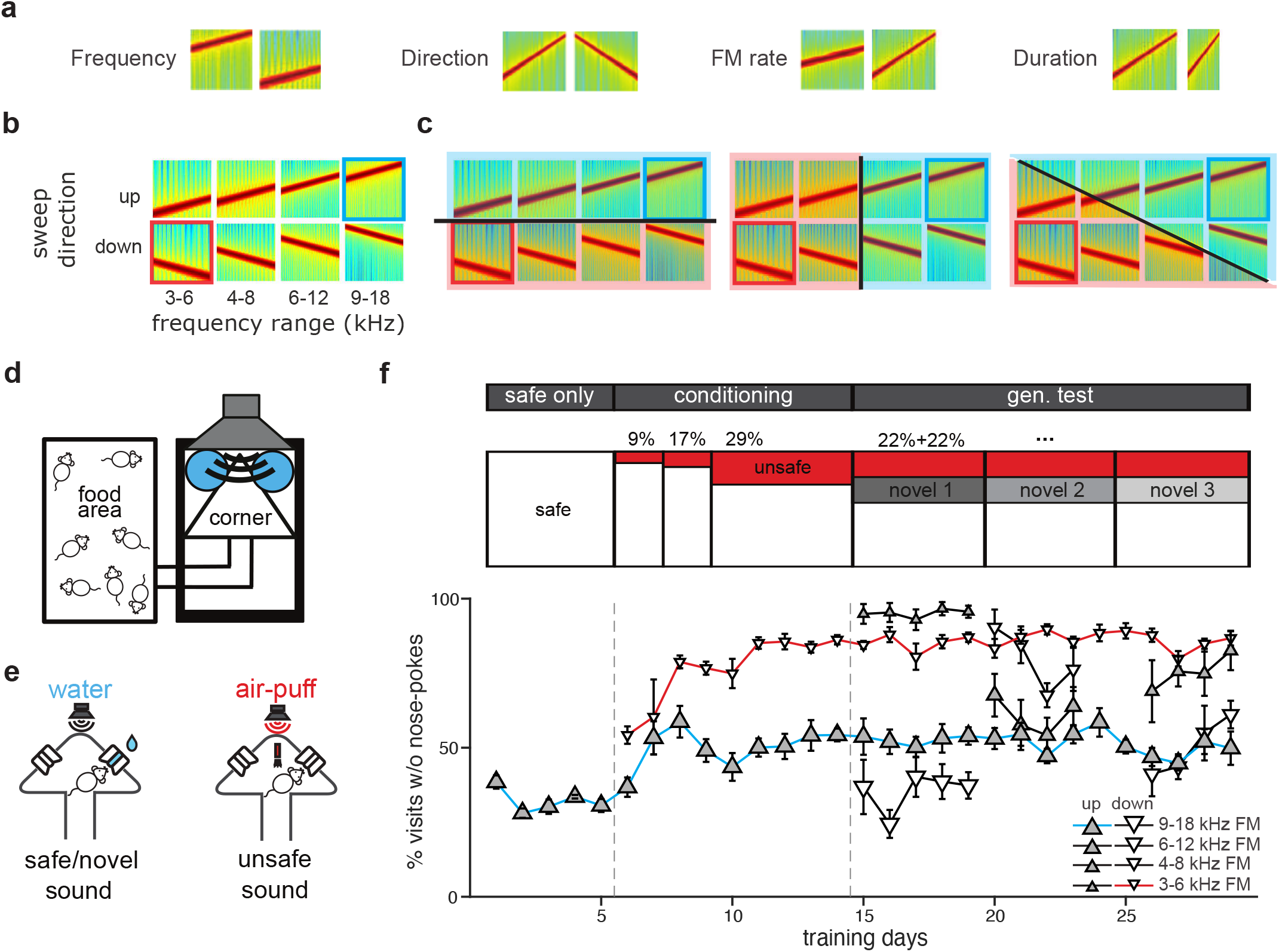
Discrimination and generalization of frequency modulated sweeps (FMs) differing in two dimensions. **a**, 4 pairs of spectrograms of FM sweeps differing in frequency range, direction, rate or duration. **b**, Spectrograms of stimulus sets varying along frequency range and sweep direction. The safe sound was a 9-18 kHz upward FM sweep (blue rectangle) and the unsafe sound was a 6-3 kHz downward FM sweep (red rectangle). **c**, Possible generalization gradients: generalisation occurring along sweep direction (left), frequency range (middle) or both dimensions (right). **d**, Schematic representation of the Audiobox. **e**, Schema of a safe/novel (left) and unsafe (right) visit. Nose-poking was followed by access to water (safe, left) or an air-puff (conditioning, right). **f**,**Top:** Experimental design. The task consists of three phases: the safe only phase with 100% safe visits (white), the conditioning phase with increasing probability of unsafe visits (red), and the generalization testing phase in which novel sounds were introduced (grey). **Bottom:** Mean daily response for mice trained with the combination of frequency range and sweep directions shown in **b**. Response was expressed as the fraction of visits without nose-pokes for different types of stimuli. Error bars represent standard error.

Each dimension pair was tested in a different group of mice. For all 4 dimensions, we chose a span of values known to elicit discriminative responses in neurons in the auditory system. The mice were trained in the Audiobox (Fig. 1d; Supplementary Fig. 1a-b), an apparatus in which mice live in groups for the duration of the experiment (several weeks) while performing the task *ad libitum*. The Audiobox consists of 2 main compartments: a home-cage where food is always available and a ‘drinking chamber’ where water is delivered upon a nose-poke. As the mouse entered the chamber, a given sound began to play repeatedly for the duration of the visit. In those visits in which the ‘safe sound’ was presented, mice had access to water (Fig. 1e, left), whereas when the ‘unsafe sound’ was played, nose-poking was followed by an aversive air puff and no access to water (Fig. 1e, right; Supplementary Fig. 1c). The task, therefore, resembled a Go/No-go paradigm. During the generalization phase, two novel sounds (appearing in 11% of the chamber visits each) were introduced, and changed every 2 to 4 days (Fig. 1f). These sounds behaved like ‘safe sounds’ in the sense that in the visits in which either of these sounds played, the door giving access to water opened upon a nose-poke (Fig. 1e, left). We measured a mouse’s response to a given sound as its level of nose-poke avoidance, i.e. the percentage chance that it will not poke its nose in that visit (the ‘response’).

### Frequency range but not direction of modulation controlled behavior

The first group of mice was trained to discriminate sweeps that differed in frequency range and sweep direction (9 to 18 kHz upward vs. 6 to 3 kHz downward sweeps, n = 9; Fig. 1b). Each day, discrimination was measured as the difference between the response to the safe sound (Fig. 1f bottom, blue line) and that to the unsafe sound (Fig. 1f bottom, red line). Mice that showed stable discrimination performance for at least three consecutive days (See Methods), were included in the analysis and then tested with novel sounds varying along both dimensions (Fig. 1b). Mice tended to either avoid or approach the novel sounds, reflecting whether they perceived them as similar to the ‘unsafe’ or the ‘safe’ sound respectively (Fig. 1f bottom, black lines).

The mean response for this first set of sounds, which varied in frequency range and modulation direction (safe, unsafe and novel, Fig. 2a, top), what we term the ‘generalization pattern’, is shown in both the form of a performance plot (bottom). We found that whether a novel sound was treated more like the unsafe or safe sound, that is how it was generalized, depended mostly on its frequency (the average correlation coefficient between frequency and behavior, ρ(freq), was -0.89) and not its sweep direction (ρ(dir) = 0.17). Regardless of the direction, mice approached novel sounds of high frequency ranges (6-12 kHz and 9-18 kHz), and avoided novel sounds of low frequency ranges (3-6 kHz and 4-8 kHz). A two-way ANCOVA revealed a significant main effect of frequency range (*F*(1,44) = 117.73 *p* = 0), and a weak effect of sweep direction (*F*(1,44) = 6.85, *p* = 0.01), but no frequency × direction interaction (*F*(1,44) = 2.52, *p* = 0.12).

**Fig. 2.**
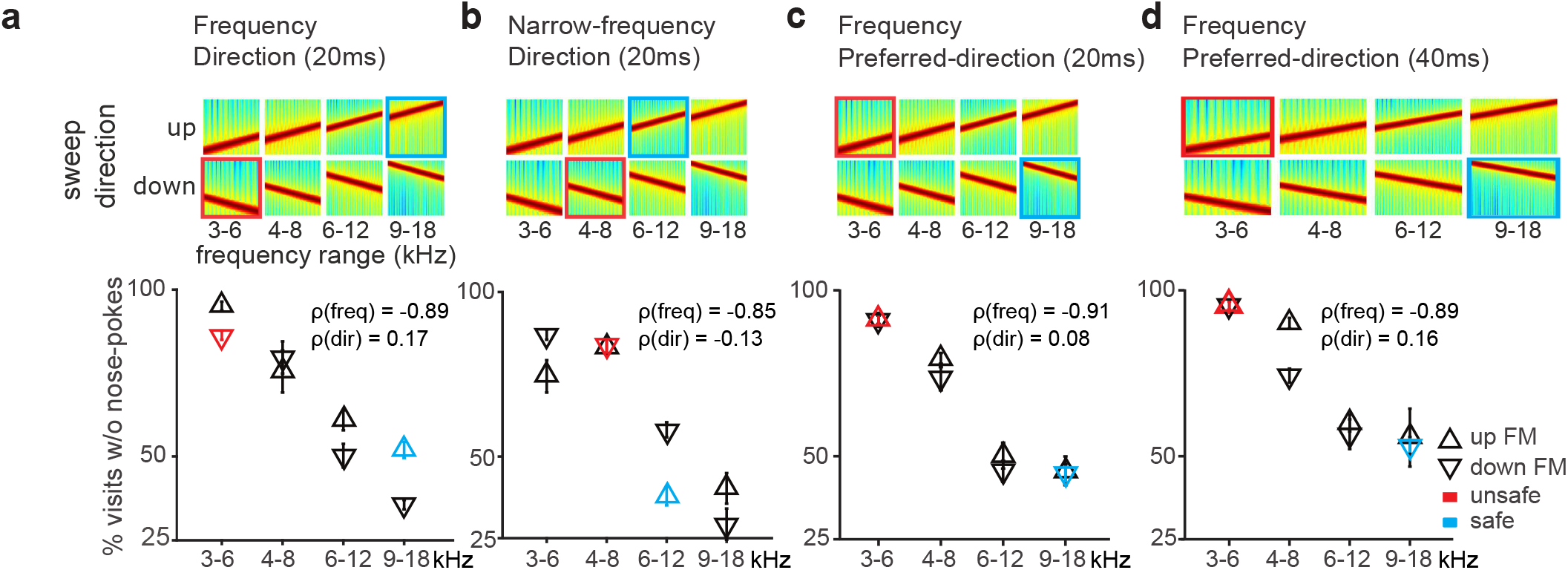
The frequency range of a FM sweep dominates over its direction. **a, Top:** Spectrograms of the stimulus set used in the task. The animals were trained to discriminate the safe sound (blue-ringed) and the unsafe sound (red-ringed) that differed in frequency range and sweep direction. They were then tested on novel stimuli (the other six sounds). **Bottom:** Generalization gradients showing the average response to each stimulus. Error bars represent standard error. **b**, Similar to **a**., for mice trained to discriminate FM sweeps with partially overlapping frequency range. **c**, Similar to **a**., for mice trained to discriminate FMs with frequency-dependent preferred direction. The sound duration used for a-c. was 20 ms, and the rate of frequency modulation was 50 octaves/s. **d**, Similar to **c**., for mice trained to discriminate FMs with a duration of 40 ms and frequency modulation at 25 octave/s.

To rule out the possibility that the dominant role of frequency resulted from a more salient difference along the frequency dimension, we narrowed the frequency dimension and trained mice to discriminate sweeps of partially overlapping frequency ranges but opposite directions (6 to 12 kHz upward vs. 8 to 4 kHz downward sweeps; *n* = 7; Fig. 2b, top). The generalization pattern revealed a similar pattern as before. Mice discriminated the two sounds well and, when exposed to novel sounds, responded mainly to their frequency range ignoring direction (the average correlation coefficient between frequency and behavior, ρ(freq), was -0.85; between direction and behavior, ρ(dir) = -0.13; Fig. 2b, bottom). A two-way ANCOVA revealed a significant main effect of frequency range (*F*(1,36) = 89.31, *p* = 0), but no effect of sweep direction (*F*(1,36) = 1.49, *p* = 0.23)) or frequency x direction interaction (*F*(1,36) = 1.62, *p* = 0.21).

Physiological studies have found a systematic representation of preferred sweep direction along the tonotopic axis at both the cortical and subcortical level, such that low-frequency neurons (with a characteristic frequency of < 8 kHz) prefer upward sweeps and high-frequency neurons prefer downward sweeps^21–23^. To further confirm that the observed frequency dominance was not caused by the particular choice of frequency-direction combination, we trained mice with sweeps of two frequency ranges that were modulated in their preferred direction (9 to 18 kHz downward vs. 6 to 3 kHz upward sweeps; *n* = 9; Fig. 2c, top). Our results showed that the mice again selectively attended to the frequency range of the novel sounds, and ignored sweep direction (the average correlation coefficient between frequency and behavior, ρ(freq), is -0.91; between direction and behavior, ρ(dir) = 0.08; Fig. 2c, bottom). A two-way ANCOVA for generalization patterns revealed a significant main effect of frequency range (*F*(1,68) = 204.86, *p* = 0), but no effect of sweep direction (*F*(1,68) = 1.04, *p* = 0.31) or frequency x direction interaction (*F*(1,68) = 0.32, *p* = 0.57) .

Up to this point we had used sweeps of 20 ms duration, based on discriminable durations in mice^24^. Since previous studies have shown that sweep direction discrimination can be limited by the sweep duration^14,17,18^, we now trained and tested mice with 40 ms long sweeps (9 to 18 kHz downward vs. 6 to 3 kHz upward sweeps; *n* = 7; Fig. 2d, top). We found that an increase in sweep duration did not change the generalization pattern (the average correlation coefficient between frequency and behavior, ρ(freq), is -0.89; between direction and behavior, ρ(dir) = 0.16; Fig. 2d, bottom). A two-way ANCOVA revealed a significant main effect of frequency range (*F*(1,52) = 122.37, *p* = 0), but no effect of sweep direction (*F*(1,52) = 3.44, *p* = 0.07) or frequency x direction interaction (*F*(1,52) = 0.54, *p* = 0.47). Taken together, these results (Fig. 2a-d) show that sound frequency is perceived independently from sweep direction and treated as behaviorally more relevant.

### The behavioral control of sweep modulation rate, direction, duration and frequency range reveals a hierarchy of behavioral relevance

To test whether and how these results extended to other dimension combinations, we performed the same experiment using sounds that differed in the combination of modulation rate and direction, sweep duration and direction, or sweep duration and frequency range (Figure 3).

**Fig. 3.**
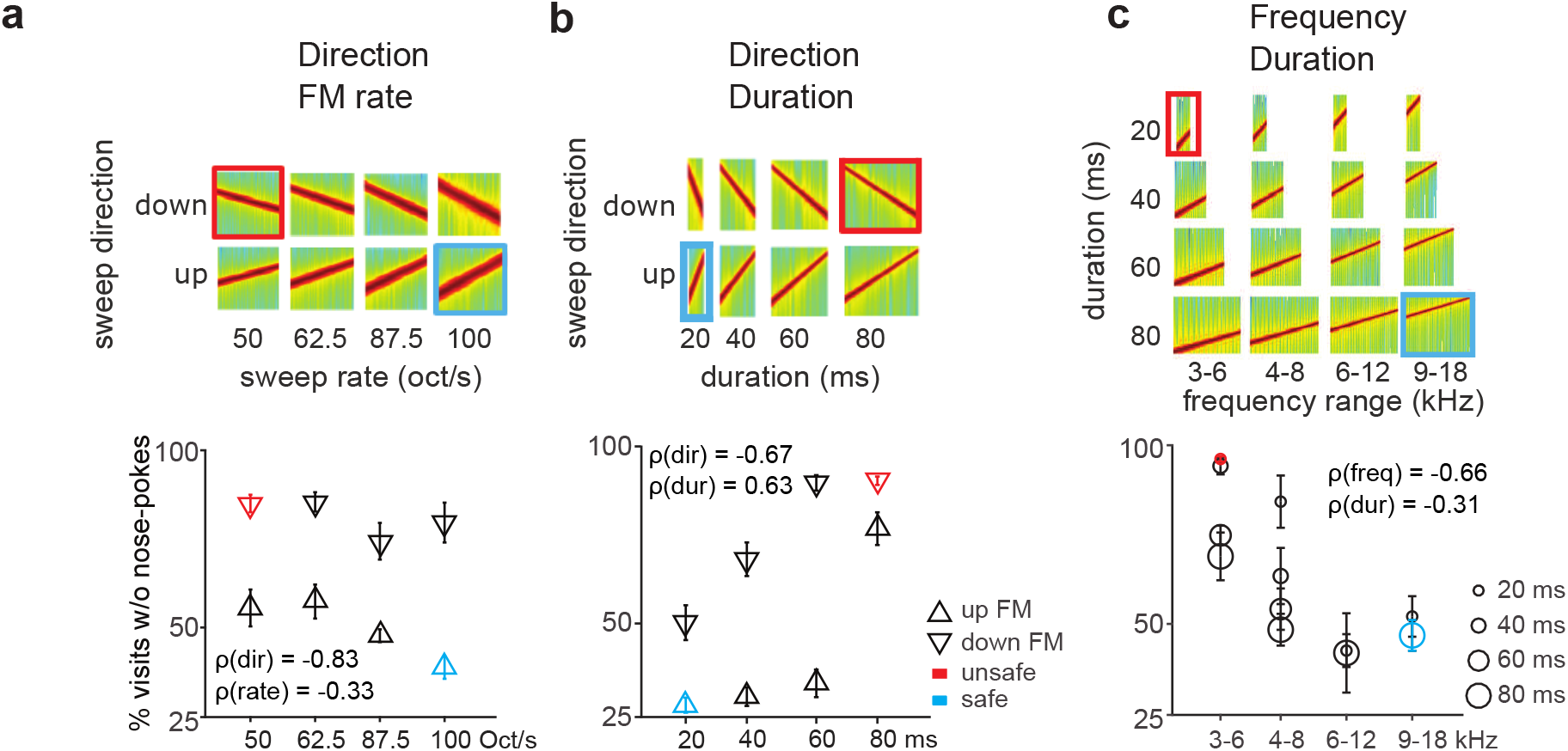
Bi-dimensional generalization reveals hierarchical perception of sound dimensions in mice. **a, Top:** Spectrograms of the stimulus set used in the task. The safe sound (blue) is a 100 octave/s upward FM sweep, and the unsafe sound (red) is a 50 octave/s downward FM sweep. All FM sweeps were centered at 8 kHz. **Bottom:** Generalization gradients showing the average response to each stimulus. Error bars represent standard error. **b**, Similar to **a**., for the combination of sweep direction and duration dimensions. Average generalization gradients (bottom) revealed an effect of both dimensions on behavior. **c**, Similar to **a**., for the combination of frequency range and duration dimensions. Average generalization gradients (bottom) showed that both dimensions controlled the animals’ behavior in a frequency-dependent manner.

We trained mice to discriminate sounds that differed in modulation rate and direction but had the same center frequency of 8 kHz (100 octave/sec upward sweep as the safe sound vs. 50 octave/sec downward sweep as the unsafe sound; *n* = 8; Fig. 3a, top). We chose 50 and 100 octave/s because these rates are within the range (25-250 octaves/s) that elicit discriminable neural responses^25^. Generalization patterns showed that mice responded to novel sounds according mainly to their sweep direction, and largely ignoring modulation rate (the average correlation coefficient between direction and behavior, ρ(dir), was -0.83; between rate and behavior, ρ(rate) = -0.33; Fig. 3a, bottom). A two-way ANCOVA analysis revealed a significant main effect of sweep direction (*F*(1,60 = 107.18, *p* = 0), a significant main effect of modulation rate (*F*(1,60) = 16.53, *p* = 0.0001), but no interaction (*F*(3,56)= 1.09, *p* = 0.30). This result further supports the idea that sound dimensions interact with each other such that a given dimension (e.g. direction) can have a stronger effect on behavior than another (e.g. rate; this experiment) but be ignored when presented with a third (e.g. frequency; previous set of experiments). This suggests a hierarchy of relevance of sound dimensions in sound discrimination.

We then tested a new set of mice with sounds that combined sweep duration and direction (20 ms 4-16 kHz upward sweeps as safe *vs* 80 ms 16-4 kHz downward sweeps as unsafe; *n* = 10; Fig. 3b, top). For the sweep duration, we used 20 and 80 ms^24^. Responses changed monotonically as a function of changes in both sound dimensions (the average correlation coefficient between direction and behavior, ρ(dir), was -0.67; between duration and behavior, ρ(dur) = 0.63; Fig. 3b, bottom). It is worth noting that changing the duration of a sweep of fixed frequency range inevitably also alters the modulation rate. Here, the modulation rates for the safe and the unsafe sound were 100 and 25 Oct/s respectively, i.e. further apart than in the previous test, where they varied between 100 and 50 Oct/s. Since in the previous experiment the role of modulation rate in discrimination was weak, it is unlikely that mice based their choice on the velocity dimension instead of the duration dimension. A two-way ANCOVA revealed a significant main effect of sound duration (*F*(1,76) = 119.23, *p* = 0), a significant main effect of sweep direction (*F*(1,76) = 132.03, *p* = 0), but no duration x direction interaction (*F*(1,76) = 0.37, *p* = 0.55). This result indicates that both dimensions contribute equally to sound discrimination.

Finally, we investigated how mice generalized along the combined dimensions of frequency range and sweep duration (9 to 18 kHz 80 ms sweeps as safe vs. 3 to 6 kHz 20 ms sweeps as unsafe; *n* = 8; Fig. 3c, top). We found that both dimensions exerted strong control over the animals’ behavior at the lower frequency ranges (around the unsafe sound) while changes in either dimension were ignored near the safe sound (the average correlation coefficient between frequency and behavior, ρ(freq), was -0.66; between duration and behavior, ρ(dur) = -0.31; Fig. 3c, bottom). A two-way ANCOVA revealed a significant main effect of frequency range (*F*(1,92) = 66.01, *p* = 0), a significant main effect of duration (*F*(1,92) = 14.13, *p* = 0.0003) and a significant interaction between the two dimensions (*F*(1,92) = 8.86, *p* = 0.004). This suggests that the influence of duration on animals’ behavior is frequency dependent.

### Mice can be trained to discriminate along the non-preferred dimension, but their learning remains localized

Up to this point we have tested the untrained response to sound stimuli varying in one or both dimensions with respect to either the safe or unsafe sounds. To further characterize the interaction dynamics between dimensions, we explored how flexible these dynamics are. With this purpose, we first trained mice to discriminate along a non-preferred dimension (e.g. sweep direction, when against frequency range) and then examined generalization again. In particular, we assessed whether the trained dimension (e.g. sweep direction, non-preferred when combined with frequency) can take over behavioral control in detriment of the other dimension (frequency range, preferred).

A new group of mice was trained to discriminate sweep pairs of opposite directions but same frequency range, so as to force direction discrimination (Fig. 4a, 5-10 kHz upward sweeps as safe vs. 10-5 kHz downward sweeps as unsafe, *n* = 8; Fig. 4a, top). Discrimination learning was slow (Supplementary Fig. 2j) but the safe and unsafe sounds were well discriminated after 50 trials of training (Fig. 4a, colored symbols). As mentioned before, direction selectivity in the auditory system is tonotopically organized, such that the physiological preferred direction of neurons of low characteristic frequencies is an upward sweep^22,23,26^. We expected, therefore, direction discrimination to be frequency-specific. Generalization to novel sounds was later measured with sweeps of successively different frequency ranges and directions (8-4 kHz downward sweeps and 6-12 kHz upward sweeps). Learning did not generalize across direction at different frequency ranges, as revealed by strong avoidance to both tested sounds despite opposite sweep direction (Fig. 4a, black symbols). To strengthen the conditioned-valence of the downward direction, we then conditioned the mice to the already tested downward sweep sound (8-4 kHz downward sweeps; Fig. 4b, colored symbols) before testing generalization again on the various novel sounds (Fig. 4b, black symbols). This conditioning did succeed in generating direction generalization across wider frequency ranges (Fig. 4b, black symbols). The average correlation coefficient between direction and behavior was -0.82, whereas the value between frequency and behavior was 0.12 (Fig. 4b). However, generalization across direction was still limited in frequency range, as reflected by the absence of categorical responses to the sweeps of frequency range beyond the trained ones (3-6 kHz upward and 18-9 kHz downward sweeps; Fig. 4b outer most black symbols). A two-way ANCOVA revealed a significant main effect of sweep direction (*F*(1,60) = 74.2, *p* = 0), but no effect of frequency range (*F*(1,60) = 1.89, *p* = 0.17) or interaction between the two dimensions (*F*(1,60) = 0.53, *p* = 0.47). Therefore, generalization along the non-preferred dimension could be trained but remained specific in frequency, possibly because plasticity occurred only within a specific region of the tonotopic map.

**Fig. 4.**
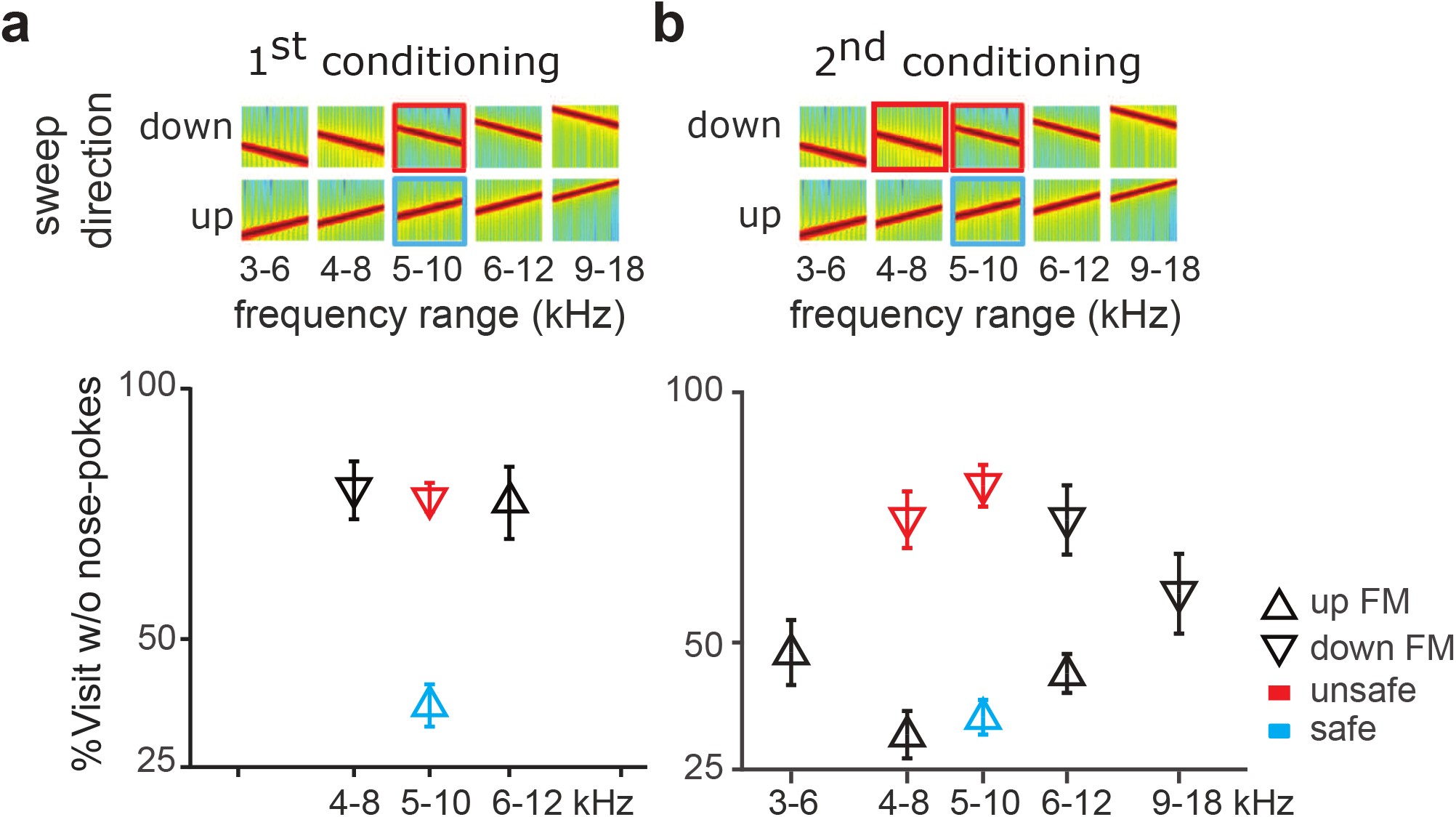
Generalization along the non-preferred dimension was localised. **a, Top**, Spectrograms of FM sweeps used in the task. The safe sound (blue) is a 5 to 10 kHz upward FM sweep, and the unsafe sound (red), a 10 to 5 kHz downward FM sweep. **a, Bottom**, Generalization gradients showing percentage of visits without nose pokes for each sound stimuli. Error bar represents standard error. **b**. Same as **a**. for generalization after the 2^nd^ conditioning during which the already tested downward FM sound (8-4 kHz downward FM sweep) was also conditioned.

### Behavioral control of sound dimensions is plastic and amenable to training

We then tested whether we could train mice that had already shown a preferred dimension to discriminate along the other, non-preferred, dimension. Here we used the animals that had been trained with the combination of frequency and sweep-direction (Fig. 2a, repeated in Fig. 5a left), where they showed a preference for the frequency dimension, while ignoring direction. These same animals were now given a 2^nd^ conditioning with the same safe sound but a high-frequency downward-sweep as the unsafe sound (Fig. 5a, middle), to encourage them to discriminate along the direction dimension. The 2^nd^ conditioning led to an increase in avoidance to the new unsafe sound (from 35% to 65%) but their discrimination did not reach levels comparable to those of the 1^st^ conditioning even after 4 days (Fig. 5a right; paired t-test for d’, *p* = 0.002). A similar result was observed in mice previously trained with the combination direction-rate (Fig. 5b, left). When the upward 50 octaves/s was explicitly conditioned to force discrimination along the velocity dimension, we observed increased avoidance to the new unsafe sound (from 56% to 69%) but weaker discrimination than in the original conditioning (Fig. 5b, middle and right; paired t-test for d’, *p* = 0.003).

**Fig. 5.**
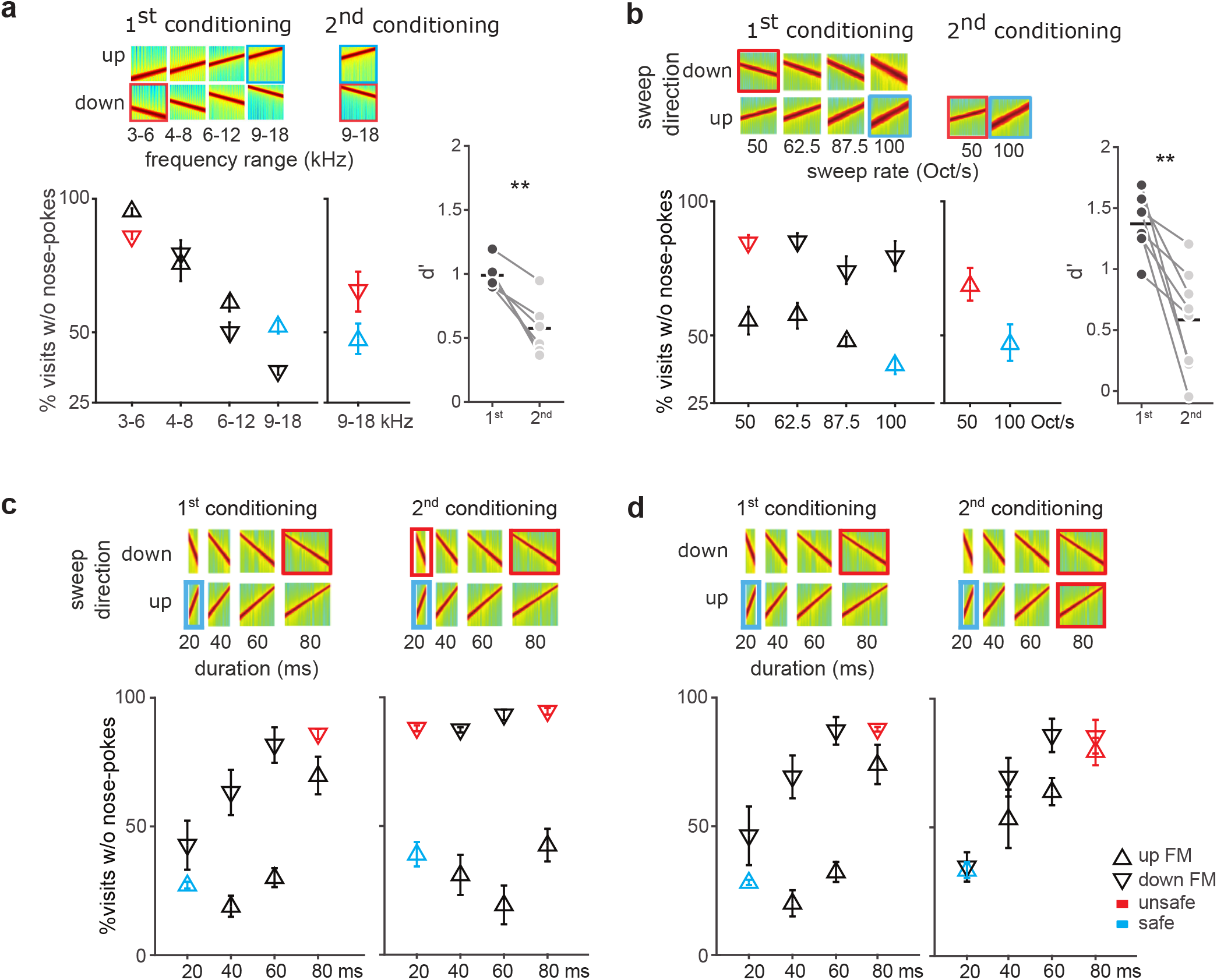
Behavioral control of sound dimensions is plastic and amenable to training. **a, Left**, Generalization gradients after initial training (1^st^ conditioning) with safe (blue square) and unsafe (red) sounds combined differences in frequency range and sweep direction. **a, Middle**, Generalization gradient after subsequent conditioning to the non- dominant dimension, sweep direction (the 2^nd^ conditioning, half the mice). **a, Right**, d’ for the last 50 unsafe visits of the 1^st^ and the 2^nd^ conditioning for individual mice. **a**, Mice initially selectively generalised along frequency range (left), and were worst at direction discrimination (middle). **b**, Same as **a**. for mice initially trained with sweep direction and rate (left), and subsequently trained to discriminate the non-dominant dimension, rate (middle). **c-d, Left**, Generalization gradients after the 1st conditioning with direction and duration. Mice initially generalised along both direction and duration dimensions. **c-d, Right**, Shift in generalization pattern after the 2nd conditioning to discriminate either direction (**c**) or duration (**d**). Responses to the safe and unsafe sound are marked as blue and red respectively.

We next examined flexibility in a task in which both tested dimensions influenced behavior. Mice that had been trained with the combination of direction-duration (Fig. 3b), were divided in two equally sized groups of comparable performances (Fig. 2c and d, left). The first half were subsequently trained with an additional unsafe sound (the 2^nd^ unsafe tone) with the same direction as the initial unsafe sound but the shorter duration of the safe sound (Fig. 5c, *n* = 5). Thus, the safe and the unsafe sounds differed now only in sweep direction. After this 2^nd^ conditioning, mice now avoided nose-poking in the presence of the added unsafe tone (Fig. 5c, right, colored symbols; 88% of avoidance, d’ = 1.45). Importantly, learning not only affected the newly unsafe sound, but was extended to novel sounds in between the safe and unsafe ones (Fig. 5c, right, black symbols). The discrimination gradient was now mainly along the direction dimension. A two-way ANCOVA for performance of the retrained mice revealed a significant main effect of sweep direction (*F*(1,36) = 264.73, *p* = 0), but no effect of duration (*F*(1,36) = 0.58, *p* = 0.45) or interaction between those two dimensions (*F*(1,36) = 0.62, *p* = 0.43).

The other half of the mice (Fig. 5d) were exposed to an additional unsafe tone that had the same duration as the first unsafe tone but the direction of the safe sound. This resulted in safe and unsafe sounds differing only in sound duration (Fig. 5d, right, colored symbols). Interestingly, even though the newly unsafe sound was already treated like an unsafe sound by the animals after the initial conditioning (from 74% to 79%; d’_2nd_ = 1.24), responses to novel sounds changed dramatically. Mice now mainly discriminated along the duration dimension (Fig. 5d, right, black symbols). A two-way ANCOVA revealed a significant main effect of sweep duration (*F*(1,28) = 51.28, *p* = 0), a weaker but significant effect of direction (F(1,28) = 6.27, p = 0.02), but no interaction between those two dimensions (*F*(3,24) = 0.65, *p* = 0.43). Thus, changes in task conditions can shift the behavioral pattern from one where both dimensions are used in discrimination to one in which one dimension is ignored. In summary, mice could be trained to flexibly use a single dimension as the behavioral relevant one, however, discrimination along non-preferred dimension remained more difficult and did not generalize beyond narrow ranges.

### Perception of ‘envelope’ features in mice

The ‘envelope’ of the sound stream is important for auditory perception, for example in speech processing^27,28^. The experiments described so far focused on differences within the short frequency-modulation bouts presented always at 3 Hz (Fig. 1-5). Next, we investigated how the ‘envelope’, i.e. the contour, influenced generalization patterns. Since amplitude modulations and the periodic envelope are important characteristics for acoustic communication signals^29^, we trained and tested mice with 100% sinusoidal amplitude-modulated (AM) stimuli (differing in dimensions of carrier frequency and modulation rate, Fig. 6a).

**Fig. 6.**
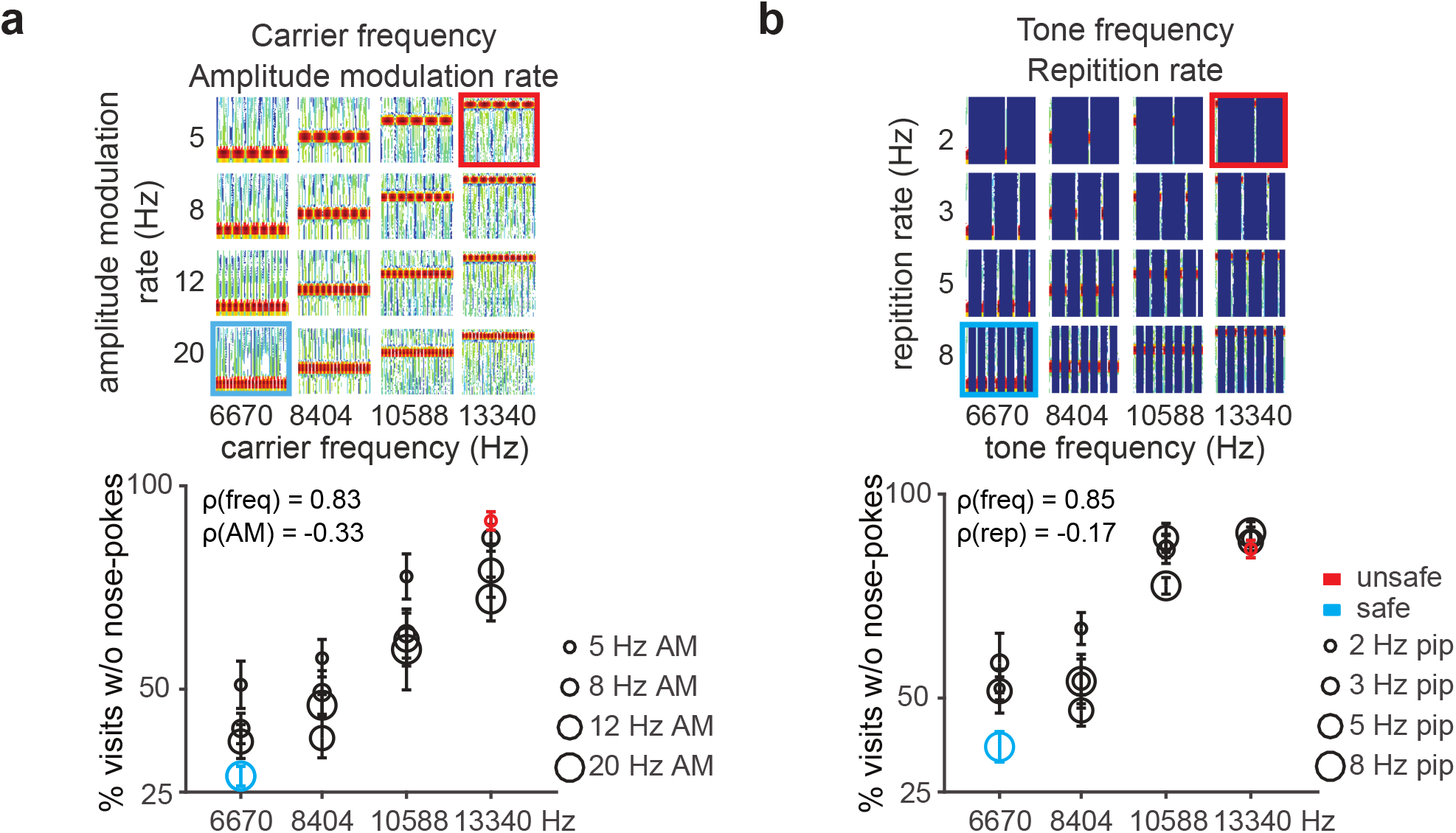
Bidirectional generalization of periodic sound. **a, Top**, Spectrograms of FM sweeps used in the task. **a, Bottom**, Generalization gradients plotting the response proportion to each of the stimuli presented in the task for the combination of tone frequency and AM sweep rate dimensions. **b**, Same as **a**. for the combination of tone frequency and repetition rate dimensions. For each figure, the safe and unsafe sounds are marked in blue and red, respectively.

We chose well discriminable carrier frequencies that differed by 1 octave^30^ and amplitude-modulations ranging between 5 and 20 Hz modulation rate^4^ (ΔF > 1 octave). Fig. 6a shows responses to trained and novel sounds. We found that, for all mice, performance was mainly controlled by the carrier frequency and influenced by the amplitude modulation (the average correlation coefficient between frequency and behavior, ρ(freq), was 0.83; between amplitude-modulation rate and behavior, ρ(AM) = -0.33). A two-way ANCOVA revealed a significant main effect of carrier frequency (*F*(1,124) = 168.76, *p* = 0), a significant main effect of modulation rate (*F*(1,124) = 26.38, *p* = 0), but no interaction between the two dimensions (*F*(1,124) = 0, *p* = 0.98).

We then tested sounds differing in dimensions of tone frequency and repetition rate (Fig. 6b) using rates ranging between 2 and 8 Hz (ΔF of 2 octaves), since neurons in the auditory cortex selectively prefer stimulus rates ranging from 2 to 15 Hz^31^. Similar to amplitude-modulated stimulus, we found that the animals’ performance was influenced by both dimensions (the average correlation coefficient between frequency and behavior, ρ(freq), was 0.85; between repetition rate and behavior, ρ(rep) = -0.17). A two-way ANCOVA revealed a significant main effect of tone frequency (*F*(1,92) = 189.3, *p* = 0), a significant effect of repetition rate (*F*(1,92) = 6.72, *p* = 0.01) and significant interaction between the two dimensions (*F*(1,92) = 4.36, *p* = 0.04). Thus, although tone frequency exerts a stronger control over behavior, repetition rate also influences behavior in a frequency-dependent manner.

### Summary

For an overview of the effect of the different dimensions on performance, we collated across tasks the correlation coefficients between each dimension tested and performance (Fig. 7a). The variability across individual animals was small, suggesting that the dimensional relevance reflects an innate neural representation in the brain. While frequency consistently exerted considerable control over the behavior across all tests, direction and duration were more variable and their contribution to behavioral decisions depended on the second sound dimension they were tested with. Temporal features such as rate of frequency modulation, amplitude modulation, or repetition rate, had less control over behavior in this type of go/no-go discrimination. Thus, the different features tested here, and in the ranges used, differed in their influence over the behavior in a way that was dependent on the other dimensions of the tested sound, revealing a hierarchy of influence dominated by frequency (Fig. 7b).

**Fig. 7.**
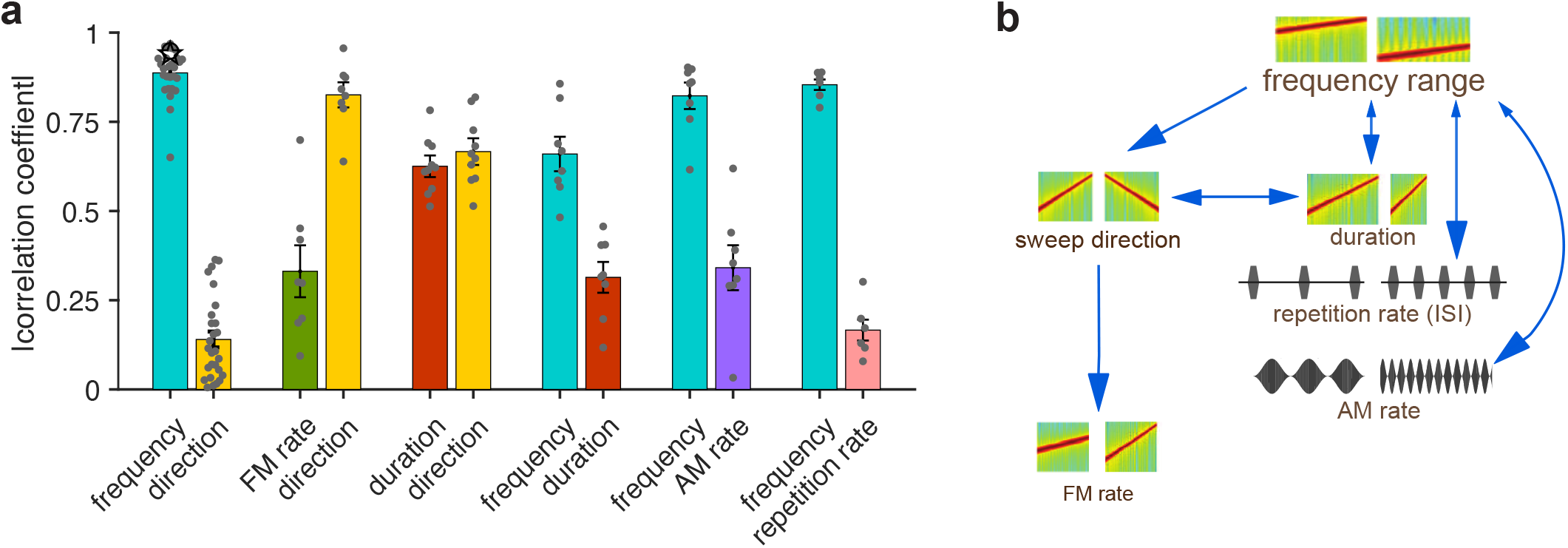
Summary. **a**, Bar plot shows the absolute value of the correlation coefficient between each dimension’s tested variable space and performance of each mouse. Error bars represent standard error. **b**, Scheme shows the perceptual hierarchy inferred by the results.

### Learning rate did not clearly influence generalization patterns

Next, we wanted to explore whether the generalization pattern was related to the learning rate during initial discrimination training between safe and unsafe sounds. With this aim, we plotted the response to the unsafe sound in blocks of 4 visits starting after the first conditioned visit (Supplementary Fig. 2 and 3, red). The average responses to the safe sound before (gray) and after conditioning (blue) are included. We did this for every dimension combination explored (Supplementary Fig. 2a-g). We also assessed learning rates for single-dimension discriminations of frequency, direction and duration (Supplementary Fig. 2h-k).

Discrimination learning in all shown frequency-direction combinations was slow (Supplementary Fig. 2a-d) when compared to discrimination learning based on frequency information alone (Supplementary Fig. 2h-i) but faster when compared to learning based on direction information alone (Supplementary Fig. 2j). For the frequency-direction combination, discrimination learning was progressively faster across tasks in the order shown in Supplementary Fig. 2a-d, and reflected in the final discrimination performance (Supplementary Fig. 2l). Despite these progressive differences in learning rate, the subsequent generalization pattern was comparable across tasks: frequency dominated over direction, and direction had a weak control over behavior throughout (Fig. 2a). For example, even though using the physiologically-preferred direction^21–23^ for the given frequency range sped up learning (compare Supplementary Fig. 2a and c), it had no effect on the role of direction in the subsequent generalization pattern.

A very different pattern of results was observed in the direction-duration combination task. When we looked at the learning rate in this task, we saw that sweep direction did not slow down learning (Supplementary Fig. 2f), despite it resulting in slow discrimination learning when alone (compared to duration; Supplementary Fig. 2k and m). One could interpret this as a lack of contribution of direction to direction-duration discrimination learning. And yet, direction contributed greatly to the generalization phase in this task.

Together, the data suggest that the learning rate does not always correlate with the influence of the learned dimensions on the perceived meaning of the sound. This pattern is also observed for the frequency-amplitude modulation and frequency-repetition rate combinations, where we observed different learning rates during initial training (Supplementary Fig. 3a-d) but similar pattern of interaction between frequency and either amplitude modulation or repetition rate in the generalization phase (Fig. 6).

### Discussion

We set out to understand how sound dimensions are integrated during auditory perception. The results have consequences for our understanding of perception and the underlying circuits. Using an automatic behavioral paradigm in mice, with a design reminiscent of the implicit bias test used in humans^32^, we tested the natural perception of combinations of sound dimensions. We trained mice to distinguish between safe and unsafe sounds that differed in two dimensions, for example frequency and sweep direction, and then tested whether they categorized the safety of novel sounds according to one or both dimensions. A complex pattern of behavioral generalization over different tasks emerged. This revealed that the perception of acoustic dimensions is hierarchically organized and that this hierarchy determines how acoustic dimensions interact with each other.

### Hierarchical organization of the acoustic dimension processing

Mice categorized novel sounds as ‘unsafe’ or ‘safe’ often on the basis of one of the two tested dimensions and ignored the other dimension. This suggests a hierarchy of relevance of the dimensions examined, such that frequency dominates behavioral decisions, followed by frequency sweep direction and duration in equal measure, while sweep rate was largely ignored (Fig. 7b). Mice can however be actively trained to discriminate along their non-preferred dimension, for example sweep direction.

Our findings are consistent with other behavioral studies that did not address the dimension question specifically. For example, rats discriminate along the frequency dimension more easily than along amplitude modulation rate^4^ or loudness^33^. In mice, maternal responsiveness to wiggling calls depends critically on both frequency and duration^34,35^, which matches our inference that frequency range and duration were at high levels of the perceptual hierarchy.

### Neurophysiological consideration of dimension perception

Is the hierarchical relevance of acoustic features reflected in the functional and anatomical organization of the auditory system? The notion that sensory systems might be organized hierarchically, i.e. into a series of stages that progressively increase abstract representations, has been a popular hypothesis for decades^36^. Hierarchical processing of sensory information has been intensively studied in the visual modality^37–40^. The influential reverse hierarchy theory proposed by Ahissar and Hochstein^41^ posits that high-level abstract inputs are prioritized for perception along the sensory hierarchy and that this hierarchy maps onto an anatomical one. The acoustic dimensions used in this study do not map well onto an anatomical hierarchy of sequential processing across different auditory stations. Neural selectivity to frequency is observed across all auditory stations, from the cochlea to the cortex, and is organized tonotopically^42^. Neural selectivity for sound duration, sweep direction, sweep rate, repetition rate, and modulation rate all appear as early as the auditory midbrain^25,26,43–45^. Thus, the observed hierarchical organization for the dimensions used in this study may not result from sequential feature extraction along different auditory stations, but from the dynamic representation of each auditory feature in an early station, probably the auditory midbrain or cortex.

The distribution of response selectivity within stations in the auditory system might nonetheless influence the functional hierarchy we observed. For example, while almost all neurons show frequency selectivity, direction or modulation rate selectivity have been found in subpopulations of neurons in auditory cortex and midbrain^22,25^. Both these dimensions are in fact somewhat embedded in the tonotopic map^22,23,26^. It is therefore plausible that a given station in the auditory system processes the direction and rate dimensions of a specific sound, albeit by neurons that are selective for a specific range of frequencies. This is consistent with our finding that the learning of non-preferred dimension discrimination is frequency-dependent, i.e. direction discrimination does not generalize well outside the trained frequency range.

How salient or relevant a given dimension is to the animal might influence the relative control this dimension has over behavior and the way sounds are generalized. On one hand, the intrinsic salience of sounds might be precisely determined by the natural statistics of the acoustic environment. It could be that frequency is always more salient than direction because it is found to be more informative in natural sounds. On the other hand, subtle differences in saliency might not affect the overall patterns of generalization and the hierarchy we observed. Although we did see differences in learning rates which might have been related with changes in saliency when we varied the characteristics of a given dimension, these did not in turn influence the generalization pattern. For example, changes in the specific direction that was paired with the low or high frequency sound influence the initial learning rate, and yet, these differences in learning rate had no real influence in the generalization pattern, which consistently showed that direction had little control over behavior compared to frequency. We were nonetheless careful to use a variable space covering a range that elicited discriminable responses in neurons in the auditory system.

To our knowledge, our experiment constitutes the first successful use of multidimensional generalization to infer untrained dimensional integration and the hierarchy of auditory information processing. These results yield important insights into the processes underlying auditory object processing and categorization. Tapping into the untrained behavioral response of mice to sounds combining different acoustic dimensions, we found that these dimensions are hierarchically organized in the control they exert over behavior. This emergent hierarchy is consistent with the predictive value of these dimensions in natural sounds as demonstrated by models. Overall, and inferring beyond the auditory modality, the data support the more general idea that both the sensitivity of physiological responses and behavioral decisions to natural stimuli has evolved to optimally use the predictive value of the different stimulus dimensions in the natural environment.

## MATERIALS AND METHODS

### Animals

A total of 103 female C57BL/6JOlaHsd (Janvier, France) mice were used. All mice were 5-6 weeks old at the beginning of the experiment. Animals were housed in groups in a temperature-controlled environment (21 ± 1°C) on a 12 h light/dark schedule (7am/7pm) with access to food and water *ad libitum*.

All animal experiments were approved by the local Animal Care and Use Committee (LAVES, Niedersächsisches Landesamt für Verbraucherschutz und Lebensmittelsicherheit, Oldenburg, Germany) in accordance with the German Animal Protection Law. Project license number 33.14-42502-04-10/0288 and 33.19-42502-04-11/0658

### Apparatus: the Audiobox

The Audiobox was a device developed for auditory research from the Intellicage (TSE, Germany). The Audiobox served both as living quarters for the mice and as their testing arena. Each animal was individually identifiable through an implanted transponder, allowing the automatic measure of specific behaviors of individual animals. The mice lived in the apparatus in groups of 7 to 10 for several days. Mice were supplied water and food *ad libitum* but are required to discriminate and respond to sounds when they go to the drinking ‘chamber’. At least one day before experimentation, each mouse was lightly anaesthetized with Avertin i.p. (0.1ml/10g) or isoflurane and a sterile transponder (PeddyMark, 12 mm × 2 mm or 8 mm × 1.4 mm ISO microchips, 0.1 gr in weight, UK) was implanted subcutaneously in the upper back. Histoacryl (B. Braun) was used to close the small hole left on the skin by the transponder injection.

The Audiobox was placed in a temperature regulated room under a 12 h dark/light schedule. The apparatus consisted of three parts: a home cage, a drinking ‘chamber’, and a long connecting corridor (Supplementary Fig. 1a, b). The home cage served as the living quarter, where the mice have access to food *ad libitum*. Water was delivered in the drinking ‘chamber’, which was positioned inside a sound-attenuated box. Presence of the mouse in the ‘chamber’, a ‘visit’, was detected by an antenna located at the entrance of the chamber. The antenna read the unique transponder carried by each mouse as it enters the chamber. The mouse identification was then used to select the correct acoustic stimulus. A heat sensor within the chamber sensed the continued presence of the mouse. Once in the ‘chamber’, specific behaviors (nose-poking and licking) could be detected through other infrared sensors. All behavioral data was logged for each mouse individually. Access to water was controlled by opening or closing of the doors behind the nose-poking ports. Air puffs were delivered through an automated valve which is placed on the ceiling of the ‘chamber’. A loudspeaker (22TAF/G, Seas Prestige) was positioned above the ‘chamber’, for the presentation of the sound stimuli. During experimentation, cages and apparatus were cleaned once a week by the experimenter.

### Sounds

Sounds were generated using Matlab (Mathworks) at a sampling rate of 48 kHz and written into computer files. Intensities were calibrated for frequencies between 1 and 18 kHz with a Brüel & Kjær (4939 ¼” free field) microphone.

Mice were trained with pairs of either frequency-modulated (FM) sweeps, amplitude-modulation (AM) sounds, or pure tone pips. For 2-dimensional discrimination task, sound pairs used in training as safe and conditioned differed in two out of four chosen dimensions. For the FM sweep sounds these dimensions were frequency range, duration, sweep direction of modulation and velocity of the sweep. For example, when the two dimensions used were the frequency range and the sweep direction, the safe sound could be an upward sweep in the high frequency range while the unsafe sound would be a downward sweep in the low frequency range. Frequency was modulated logarithmically from low to high frequencies (upward sweep) or from high to low frequencies (downward sweep). Sweeps had a duration of 20 ms (default) or 40 ms, including 5 ms rise/decay, and one of four modulation velocities (50, 62.5, 87.5 or 100 octave/sec; with 50 octaves/s as default). The sweeps were presented at roving-intensities (70 dB ± 3 dB). For the AM stimuli, tested dimensions were carrier frequency and modulation rate. The AM sounds had 100% sinusoidal modulation and had one of four carrier frequencies (6670, 8404, 10588 or 13340 Hz), as well as one of four modulation rates (5, 8, 12 or 20 Hz). For pure tone pips, the dimension of tone frequency and repetition rate were tested. Similar to the AM stimuli, the pure tones pips had one of four frequencies (6670, 8404, 10588 or 13340 Hz) and one of four repetition rates (2, 3, 5 or 8 Hz). Pure tones had a length of 20ms, with a 5ms rise/fall ramp and were presented with intensities that roved between 67 and 73 dB. For single dimensional discrimination, sounds pairs differed in sweep direction, duration or frequency were used for training.

Our choice of sounds was based on the existing literature. Rodents perceive differences in both sound frequency and sweep direction^9,12,23,25^. Mice show increased sensitivity for frequencies between 4 and 16 kHz^23^, and easily discriminate frequency distances of at least 0.20 octave^30,46^, therefore, we tested frequency ranges between 3 and 18 kHz such that the center frequency differs by more than 0.60 octave. The choice of a 50 octave/s velocity of modulation was based on both physiological^23^ and behavioral^14^ data.

The frequency-only discrimination data were from mice trained with upward frequency sweeps of the same range used in the task with 2 dimensions (3 to 6 kHz vs. 9 to 18 kHz). For direction discrimination, the mice described in Fig. 4a were used. These are mice that were trained to discriminate between sweeps of identical frequency range (5-10 kHz) but opposite direction of modulation. Data on duration-only discrimination were obtained from mice trained to discriminate FM upward sweeps (4 to 16kHz) that differed only in duration (20ms vs. 80ms).

### Discrimination task

As described in our previous study^30^, mice were trained to discriminate two sounds, the ‘safe’ and ‘unsafe sound, in the Audiobox. Throughout the duration of the experiment, one sound (i.e. 9 to 18 kHz FMs) was always ‘safe’, meaning that access to water during these visits was granted upon nose-poke. For the first 5 days, only the safe tone was played in each visit. The doors giving access to the water within the chamber were open on the first day of training and closed thereafter. A nose-poke from the mouse opened the door and allowed access to water. Another sound, a ‘unsafe one (i.e. 6 to 3 kHz FMs), was introduced in a small percentage of ‘unsafe visits and a nose-poke during these visits was associated with an air puff and no access to water (Supplementary Fig. 1c). Most of the visits, however remained safe (safe sound and access to water) as before. The probability of unsafe visits was 9.1% for the first 2 days, increased to 16.7% for the next 2 days, then stayed at 28.6% until they showed steady discrimination performance for at least 3 consecutive days (Fig. 1f).

Mice that failed to learn the task, i.e. had no differential responses to the safe and the unsafe tone for more than 2 days, were excluded from the analysis. In total, 12 out of 105 mice were excluded.

### Multidimensional generalization measurement

During generalization testing, visits consisted of 55.6% of safe visits, 22.2% of conditioned visits and 22.2% of novel visits in which a novel sound was presented. Nose poking during the presentation of the novel tone resulted in opening of the doors that gave access to water, meaning that novel visits were never accompanied by an air-puff, the were safe in nature.

Since the safe and unsafe sound differed in two dimensions, novel sounds represented all possible combinations of values used (see sounds) along each dimension. For example, when using the 9 kHz to 18 kHz upward FM sound as the safe sound and the 6 kHz to 3 kHz downward FM sound as the unsafe sound, tested stimuli (including the safe and unsafe sounds) resulted from factorially combining 4 different frequency ranges with 2 different sweep directions.

On each testing day, only two novel sounds were presented (in 11.1 % of total visits each) in addition to the safe and unsafe sounds. Each novel sound was tested for 4 days (∼50 visits). For tasks using 2 dimensions with 2 (e.g. direction) and 4 (e.g. frequency range) values respectively, the order in which novel sounds varying in either or both dimensions were tested was as shown in Supplementary Fig. 1d-e. For tasks testing 2 dimensions with 4 values each, such as the combination of frequency and duration, novel sounds were tested in the order shown in Supplementary Fig. 1f-g.

### Analysis of performance in the Audiobox

Data were analyzed using in-house scripts developed in Matlab (Mathwork). Performance traces for different stimuli were calculated by averaging the fraction of visits without nose-pokes over a 24-hour window. Discrimination performance was quantified by the standard measures from signal detection theory, the discriminability (d’). It was calculated with the assumption that the decision variables for the safe and unsafe tone have a Gaussian distribution around their corresponding means and have comparable variances. The d’ value provides the standardized separation between the mean of the signal present distribution and the signal absent distribution. It is calculated as:

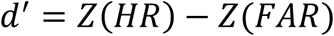

Where Z(p), pϵ[0 1], is the inverse of the cumulative Gaussian distribution, HR is the hit rate, where a hit is the correct avoidance of a nose-poke in a conditioned visit, and FAR is the false alarm rate, where a false alarm is the avoidance of a nose-poke in a safe visit. Since d’ cannot be calculated when either the false alarms reach levels of 100% or 0%, in the few cases where this happened we used 95% and 5% respectively for these calculations. This manipulation reduced d’ slightly, and therefore our d’ estimates are conservative.

### Statistical analysis

Group comparisons were made using multiple-way ANCOVAs with type I sum of squares after testing for normality distribution using the Shapiro-Wilk test. Samples that failed the normality test were compared using Wilcoxon signed rank test. Multiple comparisons were adjusted by Bonferroni correction. For analysis of data consisting of two groups we used either paired t-tests for within-subject repeated measurements or unpaired t-tests otherwise. For data consisting of more than two groups or multiple parameters we used repeated-measures ANCOVA. All multiple comparisons used critical values from a t distribution, adjusted by Bonferroni correction with an alpha level set to 0.05. Means are expressed ± SEM. Statistical significance was considered if *p* < 0.05.

