## Supplementary figures for "The perceptual categorization of multidimensional stimuli is hierarchically organized"

#### Figure S1

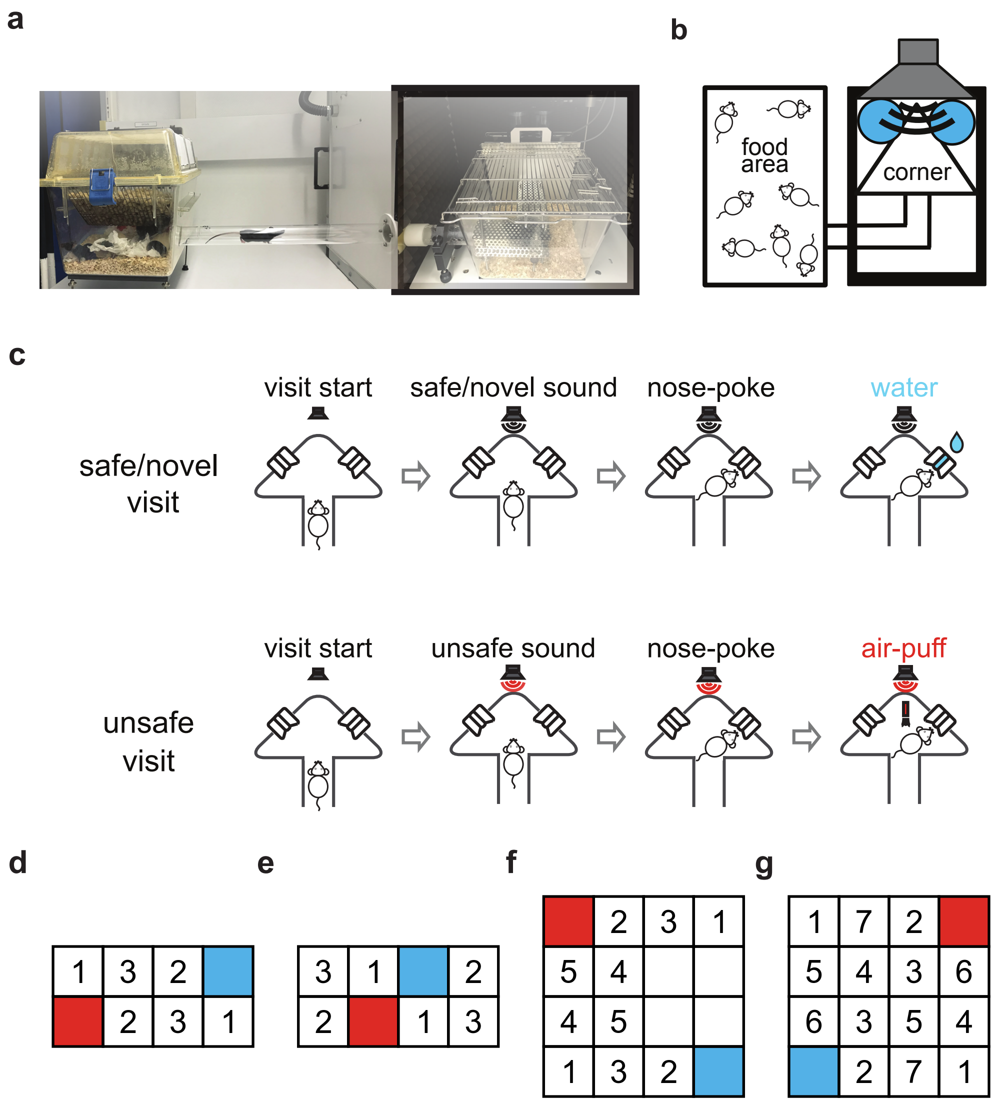


**Supplementary Fig. 1 Absolute-judgement based discrimination protocol and Audiobox apparatus.** **a**, Photos of the Audiobox. **b**, Schematic representation of the Audiobox. **c**, **Top**, Schema of a single safe/novel visit. Subjects initiated a visit by entering the corner. The sound stimulus was presented for the duration of each visit. Nose-poking was followed by access to water. **c**, **Bottom**, Schema of a single unsafe visit in which nose-poking was followed by an air-puff. **d-g**, The order of generalization testing. Blue and red rectangles represent safe and unsafe sounds. Numbers indicate the order of novel sounds being tested.

Figure S2

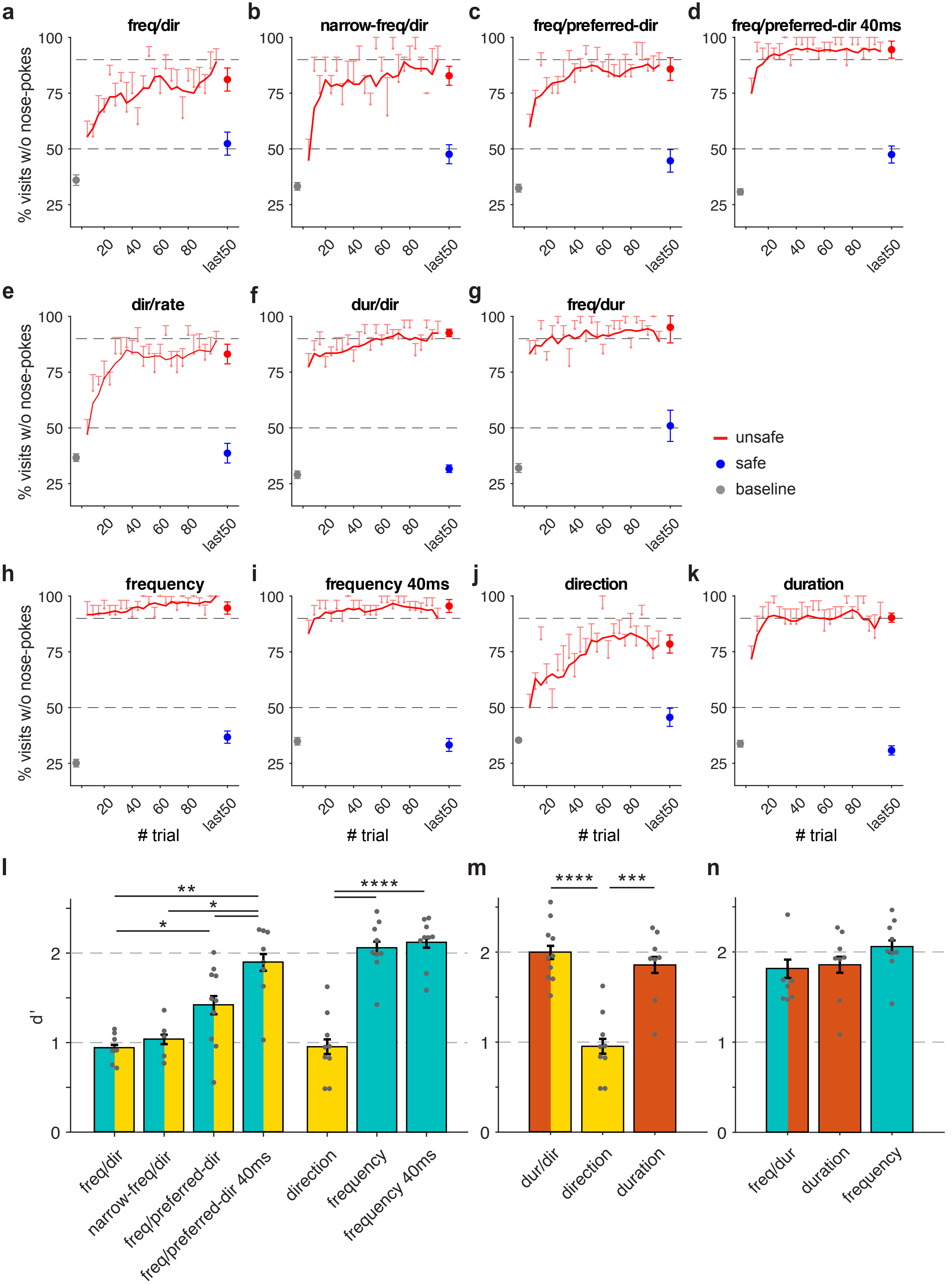
a
freq/dir
b
narrow-freq/dir
c
freq/preferred-dir
d
freq/preferred-dir 40ms
e
dir/rate
f
dur/dir
g
freq/dur
h
frequency
i
frequency 40ms
j
direction
k
duration
% visits w/o nose-pokes
### trial
last50
unsafe
safe
baseline
l
m
n
d'
freq/dir
narrow-freq/dir
freq/preferred-dir
freq/preferred-dir 40ms
direction
frequency
frequency 40ms
dur/dir
direction
duration
freq/dur
duration
frequency

**Supplementary Fig. 2 Learning curves and discrimination comparison for mice trained with FMs.**

**a-g,** Learning curves for bi-dimensional discrimination task. Responses to the unsafe sound were calculated by averaging blocks of 4 unsafe visits starting after the first unsafe visit (red). Baseline performance (grey dot) was calculated as the mean response over the last 50 safe visits before the first unsafe visit (safe-only phase). Responses to the safe (blue dot) or unsafe (red dot) sound during the last 50 unsafe visits were plotted. Learning curves were smoothed using a 5-moving average filter. **h-k**, Learning curves for discrimination along a single dimension. **l**, d' values of individual mice trained to discriminate FM sweeps differing in the frequency range (cyan) or direction (yellow) or both dimensions (cyan and yellow). **m**, d' values of individual mice trained to discriminate FMs differing in direction (yellow) or duration (red) or both dimensions (red and yellow). **n**, d' values of individual mice trained to discriminate FMs differing in duration (red) or frequency range (cyan) or both dimensions (cyan and red). Error bars represent standard error.

**Figure S3**

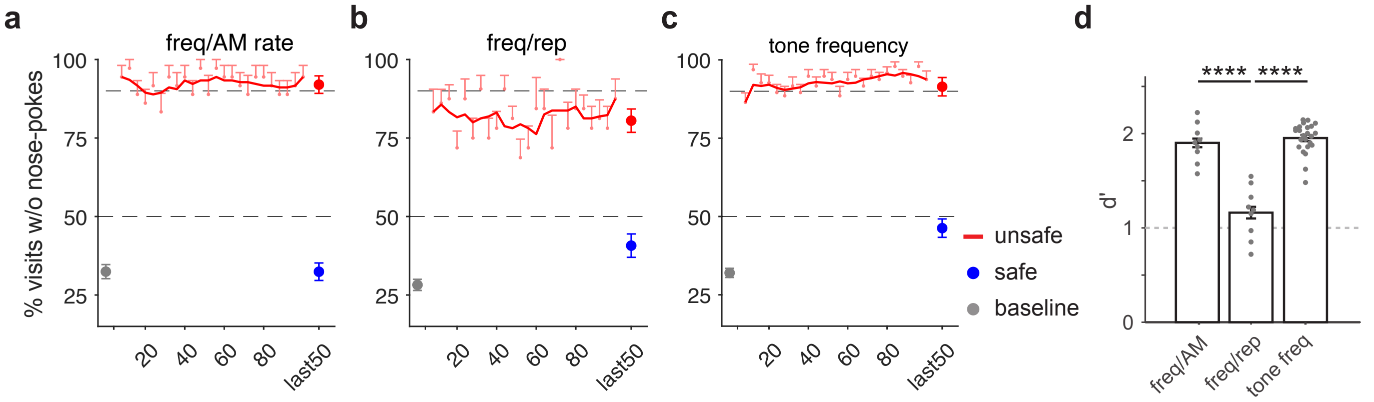


**Supplementary Fig. 3 Learning curves and discrimination comparison for mice trained with periodic sounds.**

**a-c**, Learning curves for mice trained to discriminate sounds differing carrier frequency and AM rate (**a**), tone frequency and repetition rate (**b**), or only frequency (**c**). Response to the unsafe sound was calculated by averaging blocks of 4 conditioned visits starting after the first conditioned visit (red). Baseline performance (grey dot) was calculated as the mean response over the last 50 safe visits before the first unsafe visit (safe-only phase). Average responses to the safe (blue dot) or unsafe (red dot) sounds during the last 50 unsafe visits was plotted. Learning curves were smoothed using a 5-moving average filter. **d,** d' values of individual mice trained to do uni- or bi-dimensional discrimination. Error bars represent standard error.
